# tRNA-modifying enzyme mutations induce codon-specific mistranslation and protein aggregation in yeast

**DOI:** 10.1101/2020.07.13.200121

**Authors:** Joana F Tavares, Nick K. Davis, Ana Poim, Andreia Reis, Stefanie Kellner, Inês Sousa, Ana R. Soares, Gabriela M R Moura, Peter C Dedon, Manuel A S Santos

**Author notes:** To whom correspondence should be addressed, (M.A.S.S.). These authors contributed equally to this work (Joana F Tavares, Nick K. Davis). Department Chemie, Ludwig-Maximilians-Universität München, Butenandtstr. 5-13, Haus B, B2.025, 81377 München (Stefanie Kellner).

## Abstract

Protein synthesis rate and accuracy are essential for *bona fide* protein synthesis and proteome homeostasis (proteostasis), however the mRNA translation elongation factors that prevent protein mistranslation, misfolding and aggregation are poorly understood. To address this question, we evaluated the role of 70 yeast tRNA modifying enzyme genes on protein aggregation and used mass spectrometry to identify the aggregated and mistranslated proteins. We show that the mitochondrial tRNA-modifying enzyme Slm3 thiolates the cytoplasmic tRNAs at position 34 and that decreased levels of mcm^5^s^2^U_34_ in *SLM3* mutants are compensated by increasing mcm^5^U_34_, ncm^5^U_34_ and ncm^5^Um_34_ levels. In the tRNA gene knockout strains, stress response proteins are overrepresented in protein aggregates and their genes are enriched in codons decoded by tRNAs lacking mcm^5^U_34_, mcm^5^s^2^U_34_, ncm^5^U_34_, ncm^5^Um_34_, modifications. Increased rates of amino acid misincorporation were detected in the yeast *ELP1, SLM3* and *TRM9* gene knockout mutants at protein sites that specifically mapped to the codons sites that are decoded by the hypomodified tRNAs, demonstrating that U_34_ tRNA modifications safeguard the proteome from translational errors, misfolding and cellular proteotoxic stress.

## Introduction

Transfer RNAs (tRNAs) are adapter molecules responsible for decoding mRNA into protein, playing critical roles in gene expression. In yeast, 42 different cytoplasmic tRNAs (1 initiator and 41 elongator tRNAs), encoded by 275 tRNA genes, read the 61 sense codons that code for the 20 universal amino acids (1–4). The degeneracy of this code — different codons can encode a single amino acid — is possible due to the flexibility of anticodon-codon pairing rules at the third codon position, which allows one tRNA molecule to decode more than one codon. This flexibility allows the first base of the anticodon (position 34 or the “wobble” position) to interact with more than one base at the third position of the codon (5), according to wobble-base pairing rules. To add to this complexity, tRNA nucleosides frequently undergo enzymatic modification, particularly in positions 34 and 37 of the anticodon, allowing tRNAs to differentiate between closely related codons.

The first evidence that anticodon modifications modulate the recognition of specific codons involved inosine (I) at position 34 and N^6^-isopentenyladenosine (i^6^A) at position 37 (A3’ adjacent to the anticodon) (6, 7). Unmodified U_34_ was initially believed to recognize both A and G at the third position of the codon (5). Subsequent studies revealed it could recognize all four bases at the third codon position, especially in mitochondrial translation systems (8, 9); its modification restricted recognition to only A and/or G. These findings began to functionally explain the abundance of U_34_ modifications in cytoplasmic tRNAs (10): in two-codon sets, modifications function to enhance discrimination between cognate pyrimidine-ending and noncognate purine-ending codons. This regulatory paradigm is exemplified by modifications with demonstrated roles in preventing miscoding, such as xnm^5^U in bacteria and xcm^5^s^2^U in eukaryotes (where x can be any of several different groups), as well as s^2^U, which is responsible for decoding two-codon sets that end in purine (R; NNR codons). Similarly, the presence of mcm^5^U_34_ (without thiolation) is known to improve the ability of tRNAs to read G-ending codons (11).

*In vitro* studies have also demonstrated that some tRNA modifications regulate translational accuracy by influencing the rejection of non-cognate codons at the ribosome A-site (12). Other modifications at the anticodon loop serve to stabilize the binding energetics of the codon-anticodon pair, further supporting decoding fidelity by maintaining the translational reading frame (7, 13). Meanwhile, nucleoside modifications both in the anticodon stem loop (ASL) and in the structural core of the L-shaped structure of tRNA contribute to the correct folding, stability and identity of tRNAs (7, 14–20).

Post-transcriptional modifications of tRNA have been identified in all species interrogated to date and cells commit a significant number of genes (more than 100 genes are associated with the biosynthesis of RNA modifications in yeast), as well as energy to this regulatory process. While tRNA-modifying enzymes and protein co-factors are highly conserved and appear to play important roles in regulating cellular responses to stress (21–23), most of these genes are non-essential under laboratory growth conditions (24–26), and their functions remain poorly defined. Interestingly, recent classical genetics and GWAS studies have identified single-nucleotide polymorphisms (SNPs) in genes encoding RNA-modifying enzymes in cancer, neurodegenerative diseases, diabetes, hearing loss and a range of metabolic syndromes (27), fueling speculation that these enzymes help to regulate specific subsets of disease-correlated genes. For example, mutations in the FTSJ1 methyltransferase, which modifies positions 32 and 34 of tRNA^Leu^, _^Phe^ and _^Trp^ are found in patients with non-syndromic X-linked mental retardation (28). Furthermore, the NSUN2 enzyme, which is responsible for inserting m^5^C at position 34 of tRNA^Leu^, is highly expressed in colorectal and breast cancer (29), while the MELAS and the MERRF syndromes are associated with hypomodification of mitochondrial tRNAs, namely 5-taurinomethyluridine-tRNA^Leu^ (UUR; τm^5^U) and 5-taurinomethyl-2-thiouridine-tRNA^Lys^ (τm^5^s^2^U; 30, 31). And deficiencies in 5-carbamoylmethyl and 5-methylcarboxymethyl side chains of tRNA wobble uridines (ncm^5^U and mcm^5^U, respectively) have been found to cause neurological and developmental defects in *C. elegans* (32).

While evidence suggests that mutations in RNA-modifying enzymes may contribute to the pathogenesis of serious human diseases, a limited mechanistic understanding of how these enzymes control cellular phenotype under normal physiological conditions constrains our ability to rationally evaluate these genes as potential drug targets. Here we report the development of a yeast genetic screen to identify eukaryotic tRNA modifications required for proteome stability and solubility, and whose absence results in increased protein aggregation. We used a library of 70 yeast strains, each containing a single-gene deletion of a tRNA-modifying enzyme and identified 5 genes whose absence correlated with significantly increased protein aggregation. Using homology-based spatial prediction and LC-MS/MS mapping of tRNA modifications, we found wobble uridine modifications ncm^5^U_34_, mcm^5^U_34_ and mcm^5^s^2^U_34_ to be critical for protein folding and solubility. Computational analysis of the protein aggresomes of selected yeast mutants confirmed that the absence of these modifications correlated with increased rates of amino acid misincorporation at codons decoded by hypomodified tRNAs. Together, these findings advance our understanding of the role of tRNA modifications in proteostasis, and the experimental approaches detailed herein will enable researchers to investigate previously unidentifiable consequences of RNA-related SNPs in the context of cancer and metabolic disorders.

## Materials and Methods

### Yeast library construction and growth characteristics

All *S. cerevisiae* strains used derived from BY4743 (MATa/α; his3Δ1/his3Δ1; leu2Δ0/leu2Δ0; LYS2/lys2Δ0; met15Δ0/MET15; ura3Δ0/ura3Δ0), here referred to as “wild-type” (WT). To produce mutant strains, each gene of interest was replaced by the KanMX4 cassette (EUROSCARF; Giaever et al. 2002) (Table S1). Yeast strains were grown at 30°C in YPD medium (glucose: 2% (w/v), yeast extract: 0.5% (w/v), and peptone: 1% (w/v)) (Formedium) and minimal medium without histidine (MM-His; glucose: 2% (w/v), yeast nitrogen base without amino acids: 0.67% (w/v) plus each one of the required amino acids (100 µg/ml)) (Formedium). Strains were transformed using a modified LiAc/SS Carrier DNA/PEG method (34). Briefly, cells were grown overnight in YPD at 30°C, with 180 rpm shaking, until an OD_600_ of 0.4-0.5.

Cells were then centrifuged (5000 rpm; 1 minute) and the supernatant was discarded. Transformation reagents and 3 µg of DNA (GFP-His fusion cassette amplified from the plasmid pKT128 (pFA6a–GFP(S65T)–His3MX6)) were added to the pellet and incubated at 42°C for 40 minutes. Cells were then centrifuged at maximum speed for 1 minute, the transformation mixture (supernatant) was discarded, and pellets were carefully resuspended in 200 µL of sterile milliQ water. Each cell suspension was plated on selective medium plates (MM-His) and incubated at 30°C until isolated transformant colonies were visible (2-4 days). Insertion of the fusion cassette by homologous recombination was confirmed by colony PCR for 3 biological replicates (clones).

Mutant strains were characterized by measuring growth rates as follows: (1) first, fresh medium (MM-His) was inoculated from a pre-culture and allowed to grow to an initial OD_600_ of 0.01 at 30 °C, 180 rpm, until late stationary phase; (2) next, aliquots were removed from the culture and OD_600_ was measured using a Microplate Reader (BioRad) at multiple timepoints during growth; (3) finally, time-dependent OD_600_ values were plotted.

### *In vivo* detection of protein aggregates by fluorescence microscopy

Yeast cells were harvested from 2 mL culture allowed to grow to an initial OD_600_ of 0.01 at 30 °C, 180 rpm, until early logarithmic phase, and fixed for visualization using an Axio Imager.Z1 (Zeiss) epifluorescence upright microscope, a 63X oil-immersion objective and 38 HE GFP and Brightfield filters. Images were captured using AxionVision Software (Zeiss). The images were accumulated in one representative focal plane, and subsequently processed and analyzed using ImageJ software (http://rsb.info.nih.gov/ij). Intracellular fluorescent foci were confirmed and quantified manually. On average, 500 cells of 3 clones per strain were analyzed by this approach.

### Total RNA extraction and tRNA purification

For total RNA extractions, cells were harvested from 250-mL cultures allowed to grow until logarithmic phase (OD_600_ of 1-1.5). Cell pellets were washed 3 times with ice-cold phosphate buffered saline and frozen at -80°C. Cells were resuspended in phenol-chloroform (5:1 *v*/*v*) at pH 4.7 (phenol volume = culture volume x [OD_600_/25]), and equal volume of TES buffer (10 mM Tris pH 7.5, 10 mM EDTA, 0.5% SDS). Cell suspensions were vigorously shaken for 30 sec and incubated at 65 °C for 1 h with agitation every 10 min. RNA-containing aqueous phase was separated from the phenolic phase by centrifugation at 7500 rpm for 30 min at 4°C, and re-extracted with the same volume of fresh phenol at 7500 rpm for 20 min at 4°C. Aqueous phases were once again re-extracted with the equivalent volume of chloroform-isoamyl alcohol (24:1 *v*/*v*) by centrifugation at 7500 rpm for 20 min at 4°C. RNA was precipitated overnight at -30°C with 3 volumes of pure ethanol and 0.1 volumes of 3 M sodium acetate at pH 5.2. RNA was harvested by centrifugation at 7200 rpm for 30 min at 4 °C,and resuspended in 1 mL of 0.1 M sodium acetate pH 4.5. tRNA was isolated on a 40-mL DEAE-cellulose column equilibrated with RNA resuspension buffer, as previously described (35). Briefly, samples were washed with 2.5 volumes of 0.1 M sodium acetate (pH 4.5)/0.3 M sodium chloride, and tRNA was eluted with 2 volumes of 0.1 M sodium acetate (pH 4.5)/1 M sodium chloride. tRNA was precipitated with 2.5 volumes of 100% ethanol overnight at -30°C, harvested by centrifugation and resuspended in milli-Q (mQ) water and stored at -80°C. tRNA content was subsequently verified at room temperature by electrophoresis using 15% polyacrylamide-8 M urea gels, buffered with TBE.

### Isolation of tRNA and rRNA by HPLC

Purified tRNA was analyzed using Agilent RNA 6000 pico assay before separation from rRNA by HPLC. HPLC-based purification of tRNA and rRNA (25S, 18S, 5S and 5.8S) was carried out using an Agilent 1100 HPLC series coupled to an Agilent Bio SEC-3 300 Å column (300 mm length x 7.8 mm inner diameter) with a temperature-controlled column compartment kept at 60°C and a 100-mM ammonium acetate aqueous phase (isocratic gradient) at a flow rate of 1 mL/min for 15 min. tRNA and rRNA peaks were detected by measuring absorbance at 260 nm and collected with a fraction collector using pre-determined retention times. Fractions were concentrated by SpeedVac and rehydrated with milliQ water.

### Quantitative analysis of tRNA and rRNA modifications by LC-MS/MS

Modified ribonucleosides in tRNA were identified and quantified, as reported previously (36). Briefly, isolated tRNA were enzymatically hydrolyzed and dephosphorylated by incubating for 2 h at 37°C with coformycin (50 *µ*g/mL), THU (0.3 mg/mL), MgCl2 (0.5 mM), Tris pH 8.0 (0.2 M), alkaline phosphatase (0.05 U/*µ*L), PDE1 (0.005 U/*µ*L), BHT (0.3 mM) and Benzonase (0.03 U/*µ*L). Enzymes were removed from digested ribonucleosides by ultrafiltration using a 10-kDa membrane (12000 x g, 4-10 min), and ribonucleosides were subsequently spiked with [^15^N]_5_-dA as an internal control.

Purified ribonucleoside standards were used to optimize mass spectrometry detection and fragmentation parameters. 10 pmol of each ribonucleoside standard was injected by ultra-high performance liquid chromatography (UPLC) coupled to a triple quadrupole (QQQ) mass spectrometer with 5 mM ammonium acetate as the solvent. The UPLC setup involved a Synergy 2.5-*µ*m Fusion – RP 100-Å (100 × 2 mm) column operated with a mobile phase of 0 to 80% acetonitrile (ACN) in 5 mM ammonium acetate at a flow rate of 350 *µ*L/min and 35°C. The HPLC column was coupled to an Agilent 6430 QQQ mass spectrometer with an electrospray ionization (ESI) source operated in positive ion mode with the following parameters: gas temperature = 350°C; gas flow = 10.0 L/min; nebulizer = 40.0 psi; capillary current = 5549 nA. Ribonucleoside standards were identified by HPLC retention time and collision-induced dissociation (CID) fragmentation pattern.

Digested ribonucleosides were analyzed by UPLC-MS/MS, and modifications were quantified by dynamic multiple reaction monitoring (MRM) using molecular transitions previously calculated from ribonucleoside standards. After background subtraction, the signal intensity of each ribonucleoside was normalized against signal intensity of the [^15^N]_5_-dA internal standard, permitting adjustments for day-to-day fluctuations in MS sensitivity. Signal intensity for each ribonucleoside was also normalized by dividing the raw peak area for the ribonucleoside by the sum of the UV absorbance peak areas for the four canonical ribonucleosides, in order to adjust for variations in total tRNA content in each sample. For each sample, measurements were acquired across 3 biological replicates (clonal isolates) using 3 technical replicates, and statistical significance was determined using a Student’s *t*-test. Quantified nucleoside modification data for each mutant strain was subsequently transformed to log_2_(fold change) ratios relative to WT. Hierarchical clustering analysis was performed using Cluster 3.0: average linkage algorithm based on the distance between each data set measured using Euclidean distance, with the heat map representations produced using Java TreeView.

### tRNA isoacceptor quantification by four-leaf clover qRT-PCR

Hypomodified tRNA was quantified using a polymerase chain reaction- (PCR-) based method described in Honda et al., 2015 (37). Sequences of adapters and primers for four-leaf clover reverse transcriptase PCR (FL-qRT-PCR) are shown in Table S2. Briefly, total RNA was subjected to deacylation treatment (incubation in 20 mM Tris-HCl pH 9.0 at 37 °C for 40 min), followed by annealing and ligation with a DNA/RNA-hybrid stem-loop adapter using T4 RNA ligase 2 (New England Biolabs), which specifically ligates the adapter to mature tRNA. Ligated tRNAs were specifically amplified and quantified by TaqMan qRT-PCR with an Applied Biosystems 7500 Real-Time PCR System, using a standard program of 95 °C for 20 sec followed by 40 cycles (95 °C for 5 sec, Tm for 34 sec). All reactions were run in triplicate and the threshold cycles (Ct) were determined. 5S rRNA was quantified for use as an internal control with SsoFast EvaGreen Supermix (BioRad). The amplified cDNA was developed by 10% native PAGE. For statistical evaluations of the determined CP and relative expression variations, data were analyzed for significant differences by REST-MCS© using 3 independent clones with 3 technical replicates (38).

### Extraction of insoluble aggregates by differential centrifugation

Insoluble protein fractions were isolated as described by Koplin et al., 2010, with a few modifications (39). 20 OD_600_ units of logarithmically growing cells cultivated in MM-His were harvested at 4000 rpm for 10 minutes at 4°C, and resulting cell pellets were washed with ice-cold phosphate-buffered saline (PBS) and frozen at -80°C. To prepare cell lysates, pellets were resuspended in 500 *µ*L of lysis buffer (20 mM Na-phosphate, pH 6.8, 10 mM DTT, 1 mM EDTA, 0.1% (*v*/*v*) Tween, 1 mM PMSF, protease inhibitor cocktail (Roche), 3 mg/mL lyticase and 1.25 U/mL benzonase), and incubated for 30 min at 30°C. Glass beads were used to disrupt yeast cells using a Precellys(tm) 24 disrupter; 2 cycles of 25 sec at 6500 rpm; samples were kept on ice between each cycle. Cell lysates were then centrifuged for 20 min at 200 x g at 4°C to isolate supernatant protein fractions. Following centrifugation, supernatant fractions were aspirated and adjusted to the same protein concentration (5 mg/mL for protein gels) across samples. Following aspiration of the supernatant, membrane and aggregated proteins were isolated by centrifugation at 16000 x g for 20 min at 4°C. Following this round of centrifugation, resulting supernatant fractions were aspirated and membrane proteins were removed by resuspending aggregated proteins in 2% NP-40 (in 20 mM Na-phosphate, pH 6.8, 1 mM PMSF and protease inhibitor cocktail), disrupting the mixture by probe sonication (3 × 5 sec at cycle 0.1 and amplitude 20%), and centrifuging the mixture at 16000 x g for 20 min at 4 °C. This process was repeated twice, after which final insoluble protein fractions were washed with NP-40-deficient buffer (centrifuging the mixture at 16000 x g for 20 min at 4 °C), solubilized in 50 µL of Urea Buffer (50 mM Tris-HCl, pH 7.5, 6 M urea and 5% SDS), boiled in 1X laemli sample buffer, separated by SDS-PAGE (14%) and resolved by Coomassie staining.

For Mass spectrometry analysis, insoluble protein aggregates dissolved in NP-40-deficient buffer were precipitated with TCA (100% *w*/*v*; 1 vol of TCA to 4 vol of protein sample) at 4°C for 30 min. Precipitated protein was washed with 200 *µ*L ice cold acetone (9:1 *v*/*v*), and pellets were allowed to air dry at RT. Dried protein pellets were resuspended in 50 *µ*L of 8 M urea before further processing for MS/MS analysis.

### Pan-proteome quantification of amino acid misincorporations

Insoluble protein was reduced with 10 mM DTT (1 h at 56°C) and alkylated with 55 mM iodoacetamide (1 h at RT in the dark). Protein was then digested overnight using modified trypsin (Thermo Scientific) at an enzyme/substrate ratio of 1:50 in 100 mM ammonium acetate (pH 8) at 30°C. Synthetic peptides mimicking amino acid misincorporations (designed in house) were added as internal controls to each sample (Table S3). Peptide mixtures were desalted using MicroSpin C18 columns (The Nest Group, Inc) and analyzed using an Orbitrap Fusion Lumos mass spectrometer (Thermo Scientific, San Jose, CA, USA) coupled to an Easy-nLC™ 1200 liquid chromatography system (Thermo Scientific (Proxeon), Odense, Denmark). Peptides were loaded directly onto the analytical column and separated by reverse-phase chromatography using a 50-cm column with an inner diameter of 75 μm, packed with 2 μm C18 particles (Thermo Scientific, San Jose, CA, USA). A 90-minute chromatographic gradient using 0.1% formic acid in water (buffer A) and 0.1% formic acid in ACN (buffer B) was performed for each sample analysis at a flow rate of 300 nL/min as follows: 0-79 min = start at 5% buffer B and gradually increase to 78% buffer B; 79-90 min = gradually decrease to 35% buffer B. The column was washed for 10 min using 95% buffer B after each analytical gradient, and stored overnight at 95% buffer A.

The Orbitrap was operated in data dependent acquisition (DDA) mode and full MS scans (with 1 micro scan at resolution of 120,000) were used over a mass range of 350-1,500 m/z. Auto gain control (AGC) was set to 1E5 and dynamic exclusion to 50 sec. In each cycle of DDA analysis, ions that met defined detection criteria (peptide isotopic profiles, charge states between 2+ and 5+, ion count above 1E4) were selected for fragmentation at normalized collision energy of 28%. Fragment ion spectra produced via higher-energy collision dissociation (HCD) were acquired using the following Orbitrap settings: AGC = 3E4, isolation window = 1.6 m/z, and maximum injection time = 80 ms. All data was acquired and pre-filtered using Xcalibur (v3.0.63).

MS/MS raw data was analyzed using PEAKS Studio (v.8.0, Bioinformatics Solutions Inc.) for peptide identification and label-free quantification: (1) Samples were searched against a *S. cerevisiae* database sourced from the *Saccharomyces* Genome Database (version of July 2017), which included a comprehensive list of decoy entries and common contaminants; (2) trypsin was designated as the digestive enzyme and a maximum of three missed cleavages were allowed; (3) carbamidomethylation was set as a fixed modification and oxidation (M) was set as a variable modification; (4) searches were performed using mass tolerances of 7 ppm for MS scans and 20 mmu for MS/MS scans; and (5) resulting search results were filtered for FDR ≤ 1%. Differential analysis of protein abundance was analyzed with DAPAR and ProStaR packages of R. Samples were normalized with the quantile centering method, setting the value of quantile to 0.15%, assuming that the signal/noise ratio is roughly the same in all the samples. Missing values in the 3 biological replicates of a mutated strain were imputed to value 1, whereas missing values in 1 or 2 biological replicates were imputed to the minimum value of the respective sample. Upregulated aggregated proteins (2-logFC) were identified using a Limma statistical test, with FDR ≤ 5% and p value ≤ 0.05 after calibration (pounds).

Amino acid misincorporations were identified using the SPIDER algorithm of PEAKS Studio (v.8.5, Bioinformatics Solutions Inc.), as described above with several changes: (1) raw data was searched against the *S. cerevisiae* database; (2) trypsin was chosen as the digestive enzyme with a maximum of three allowable missed cleavages; (3) carbamidomethylation was set as a fixed modification, whereas oxidation (M), Asn→Lys substitution, Asp→Glu substitution, His→Gln substitution, Phe→Leu substitution, and Ser→Arg substitution were set as variable modifications; (4) searches were performed using mass tolerances of 7 ppm for MS scans and 20 mmu for MS/MS scans; (5) additional search parameters were analyzed to identify common and unspecified post-transcriptional modifications (PTM); and (6) resulting search results were filtered for FDR ≤ 1%. Amino acid misincorporations were validated bioinformatically by programming misincorporations into a new *S. cerevisiae* protein database and re-searching raw data with PEAKS without designating amino acid substitutions as variable modifications. Amino acid misincorporations were considered valid only if bioinformatically mutated proteins (encoding mutant peptide sequences) were accurately identified in the second database search at ≤ 1% FDR.

## Results

### Deletion of tRNA modification genes induces protein aggregation

*S. cerevisiae* encodes 73 genes involved in tRNA modification, of which 21 have been shown to be essential or affect growth. We engineered a diploid yeast gene knockout library to characterize protein aggregation phenotypes of 50 homozygotic nonessential genes (two mutants – Dus1Δ and Pcc1Δ failed quality control), as well as 17 heterozygotic (single-copy deletions) essential genes or genes that confer significantly retarded growth (Tables S1 and S4). Our knockout library included the catalytic subunits of tRNA-modifying enzymes, as well as other enzymes that indirectly affect tRNA modifications — such as Kti12, a kinase required for Elongator function *in vivo*.

We developed a microwell plate-based fluorescence aggregometry platform to assess how shifts in endogenous tRNA modification profiles affect protein aggregation phenotypes in yeast. This assay employs a protein aggregate sensor we engineered by fusing Hsp104 and GFP open reading frames under control of the yeast Hsp104 promoter (Figure 1A). As Hsp104 is a disaggregase known to recognize and bind aggregated proteins, our Hsp104-GFP construct permitted rapid visualization and quantification of protein aggregates by epifluorescence and confocal microscopy (40, 41). This sensor was transformed into each strain in our KO library, and resulting cells were screened by epifluorescence microscopy (Figure 1B). Cells containing highly soluble protein content had GFP fluorescence homogenously distributed throughout the cytoplasm, while cells containing protein aggregates displayed highly concentrated fluorescent loci (Figure 1C *left*). We further analyzed these strains by transmission electron microscopy (TEM), to distinguish protein aggregates from organelle membranes also present in the cytoplasm (Figure 1C *right*, red arrows). Using TEM to analyze WT cells, we did not detect the electron-dense aggregates observed in mutant strains (Figure 1C).

**Figure 1.**
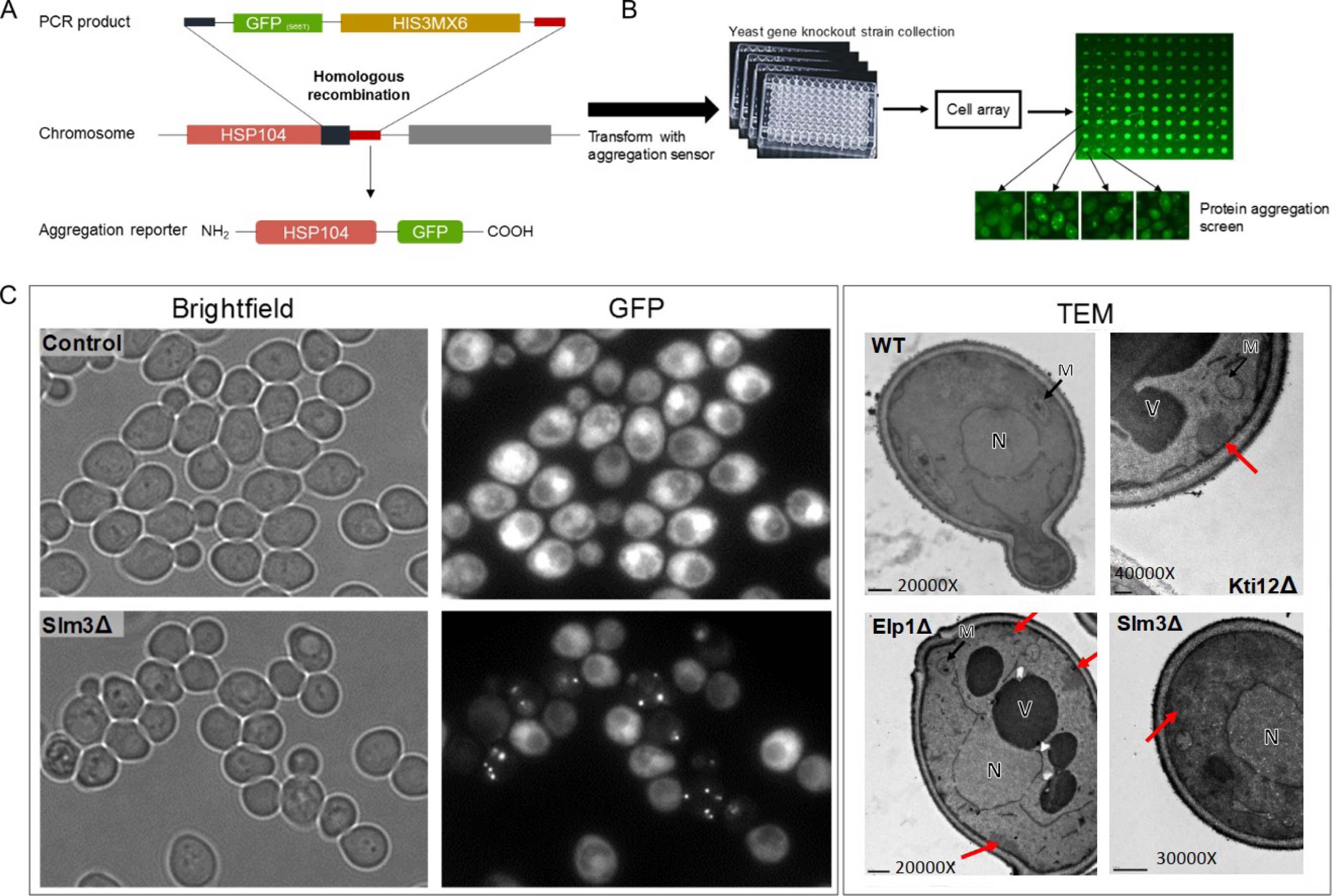
Screen to identify RNA-modifying enzyme KOs that induce protein aggregation. **A**. Engineering scheme for the construction of a fluorescent molecular sensor of protein aggregation. PCR products containing a GFP tag and a selectable marker gene (plasmid pKT128 – pFA6a-GFP(S65T)-His3MX6) were fused in-frame to the C-terminal coding region of the Hsp104 gene through homologous recombination, yielding an Hsp104-GFP fusion protein. Adapted from (73). **B**. Diagram outlining experimental strategy of screening our engineered fluorescent library to identify RNA-modifying enzyme KOs with protein aggregate phenotypes. **C. *Left box*:** WT (BY4743) and the mutant Slm3Δ yeast strains expressing the Hsp104-GFP reporter protein were collected in mid-exponential growth phase and observed by fluorescence microscopy (60x objective). Cells harboring deletion in the Slm3 gene showed localized Hsp104-GFP fluorescence, indicating the presence of protein aggregates. ***Right box*:** Ultrastructure of WT and Elp1Δ, Kti12Δ and Slm3Δ mutant yeast cells captured using Transmission Electron Microscopy (TEM) of Epon 812 resin embedded cells. Red arrows indicate regions with increased electron-dense protein aggregate content. N: nucleus; M: mitochondrion; V: vacuole.

Relative to WT, cells with deletion of Elp1, Elp3, Kti12, Slm3 and Trm9, demonstrated significantly increased localization of Hsp104-GFP into aggregate foci (Figure 2). Notably, these five strains harbor deletions in genes encoding enzymes that are involved in modifying tRNA wobble uridines. Trm9 is a wobble uridine methyltransferase that catalyzes the esterification of modified uridine nucleosides, resulting in the formation of mcm^5^U_34_ and mcm^5^s^2^U_34_ in specific tRNAs (42). Elp1 and Elp3 are one of the subunits in the Elongator complex, which is required for the formation of the cm^5^U side chain in some wobble uridines (43). Kti12 is important for the phosphorylation of Elp1 and is also required for the formation of mcm^5^ and ncm^5^ side chains at some wobble uridines (43). In contrast, Slm3 is known as a mitochondrial tRNA-specific 2-thiouridylase 1 (Mtu1), which is responsible for the formation of the s^2^ group in cmnm^5^s^2^U_34_-containing tRNA species (only in mitochondrial tRNAs) (31). Of note, the respective formation of mcm^5^ and s^2^ groups occurs through independent biosynthetic processes.

**Figure 2.**
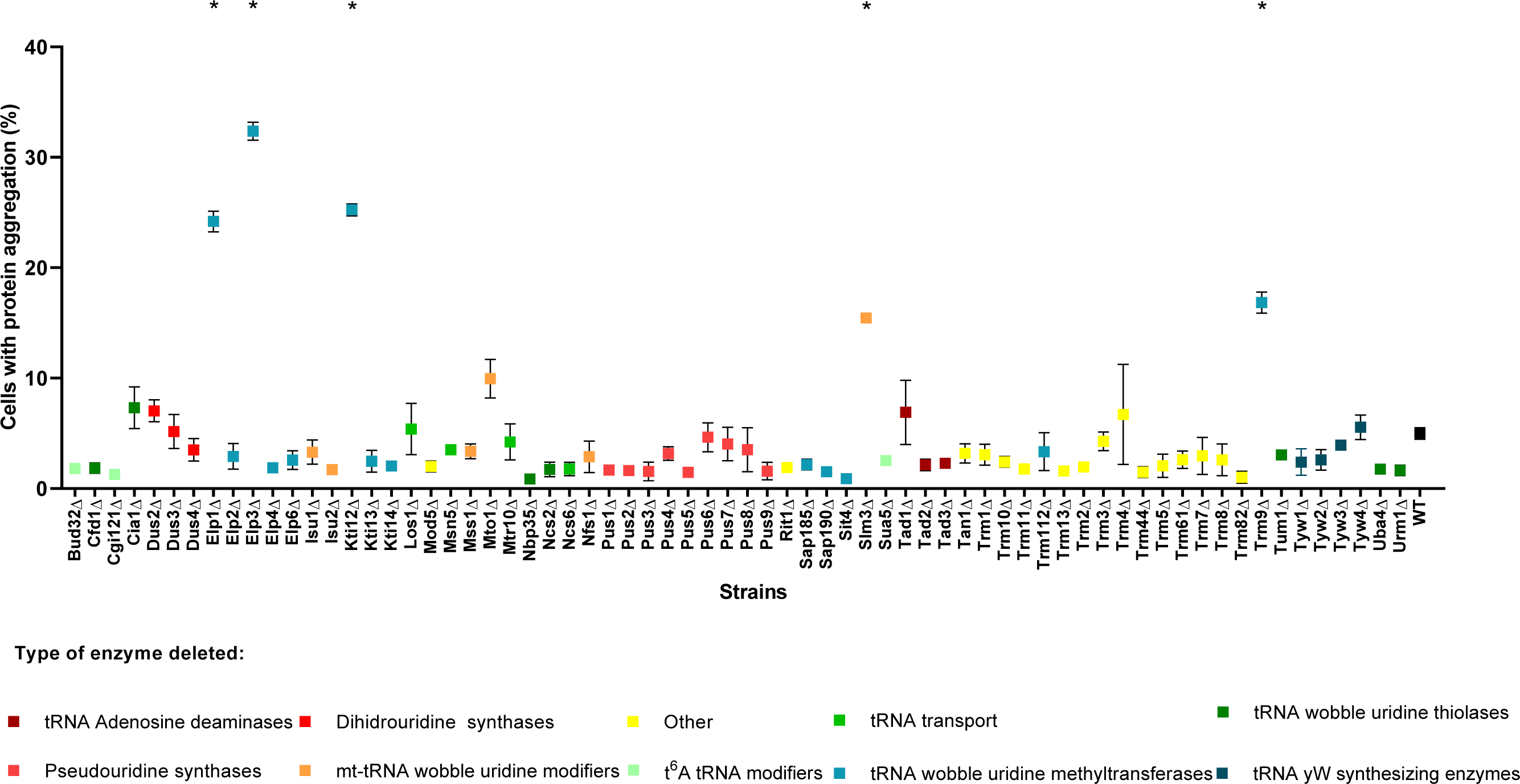
Protein aggregate phenotypes vary amongst tRNA-modifying enzyme KO strains. Data shows the mean ± SEM of the percentage of cells containing localized Hsp104-GFP fluorescent foci. For each KO strain, data represents triplicates of 3 independent clones (*p<0.001 One way ANOVA post Dunnett’s multiple comparison test and CI 95% relative to WT).

We next examined whether KO strains with significantly increased protein aggregation also exhibited slower growth and/or loss of cell viability (Figure 3), and observed a strong decrease in the cells growth relative to WT in the Elp1Δ, Kti12Δ and Trm9Δ strains, which agreed with previous reports (44–46). However, the other KO strains also exhibited slower growth rates relative to WT (Figure 3A, B). Cell viability appears not to be significantly altered (Figure 3C). Together, these results indicate that cytotoxic protein aggregates result from the disruption of certain tRNA-modification genes, and both differentially retard cell growth and promote cell death.

**Figure 3.**
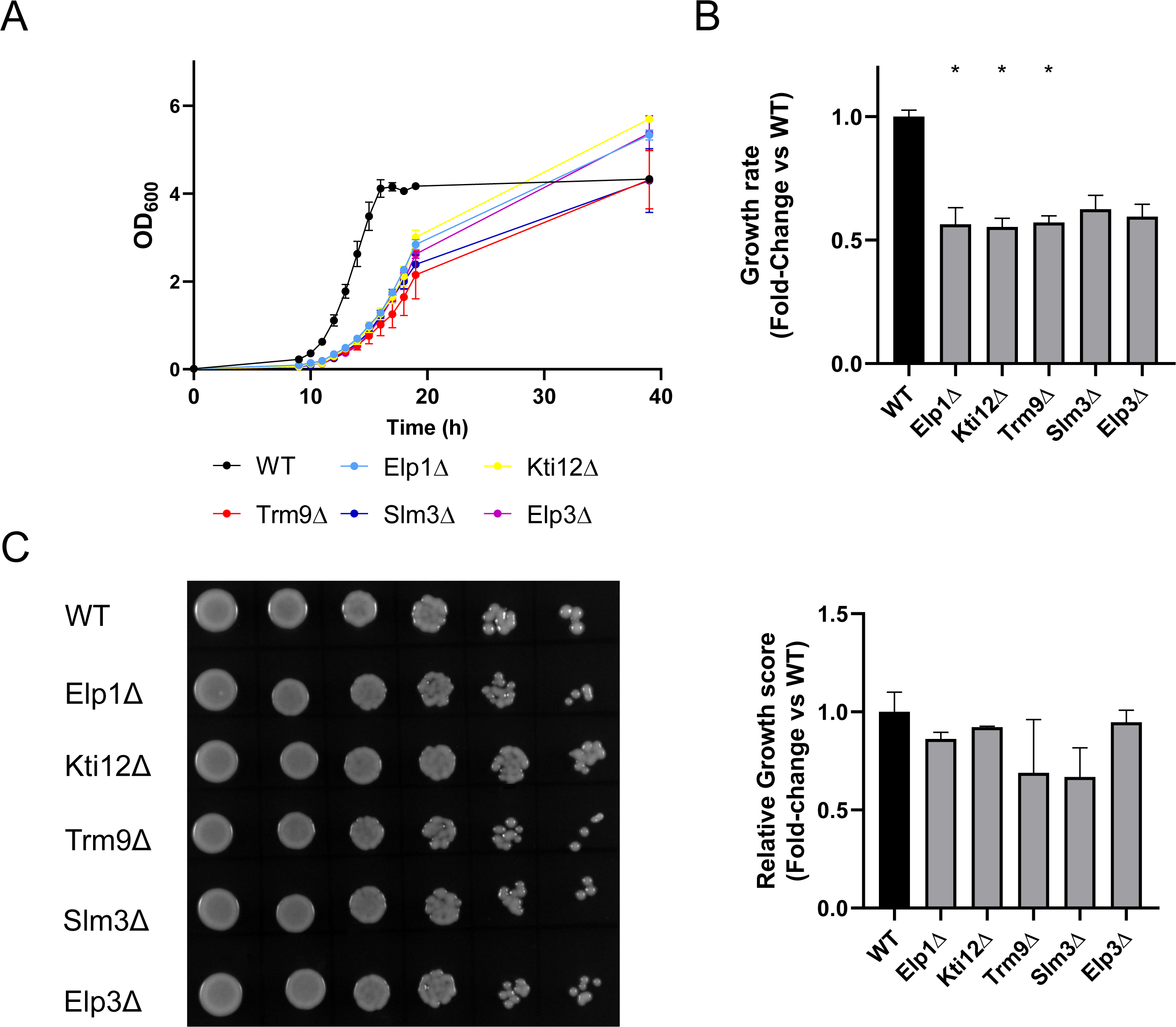
Physiological characteristics of selected RNA-modifying yeast mutants. **A**. Yeast cultures were inoculated at an initial OD_600_ of 0.01 and grown to stationary phase in selective medium (MM-His) at 30 °C and 180 rpm. Optical density was measured at timepoints along the growth curve. **B**. Relative growth rate of mutant strains was determined by normalizing the exponential growth rate of each KO strain to the exponential growth rate of WT cells. Data represents the mean ± SD of duplicate measurements for each of three independent clones (*p<0.05 Kruskal-Wallis test post Dunn’s multiple comparisons test with Benjamini-Hochberg adjustment and CI of 95%, relative to WT). **C**. Colony survival of selected mutant strains compared to WT strain. Yeast cells grown in stationary phase were diluted to an initial OD_600_ and 6 fivehold serial dilutions onto MM-His plates were spotted. Their colony-forming abilities were then analyzed after 2 days of incubation at 30 °C. The relative growth score represents a ratio between growth of each KO strain normalized against growth of the WT strain. Data represent mean ± SD of three independent clones.

### Deleting tRNA modification genes alters tRNA stability and modification levels

RNA is modified by a naturally occurring system of chemical modifications, some generated through related biosynthetic pathways, that coordinately mediate cellular physiology (7). We used LC-MS/MS to quantitatively profile variations in the global abundance of tRNA modifications across 4 of the 5 tRNA-modifying enzyme mutants selected by fluorescence aggregometry (Figure 4A). Elp3Δ was not analyzed as it was already described in the literature (47, 48). Analysis by LC-MS/MS revealed decreased ncm^5^U and ncm^5^Um levels that dropped to limits close to detection in Elp1Δ and Kti12Δ, while mcm^5^U and mcm^5^s^2^U levels were also significantly reduced in Trm9Δ, Elp1Δ and Kti12Δ (Figures 4B and S2; Table S5). Meanwhile, the Slm3 thiolase KO demonstrated both decreased mcm^5^s^2^U and increased mcm^5^U, which is consistent with the absence of uridine thiolation (Figures 4B and S2; Table S5). We also measured statistically decreases in yW levels across Kti12Δ, Trm9Δ and Slm3Δ. These findings confirmed that inactivation or deletion of tRNA-modifying enzymes concomitantly alters levels of related tRNA modifications.

**Figure 4.**
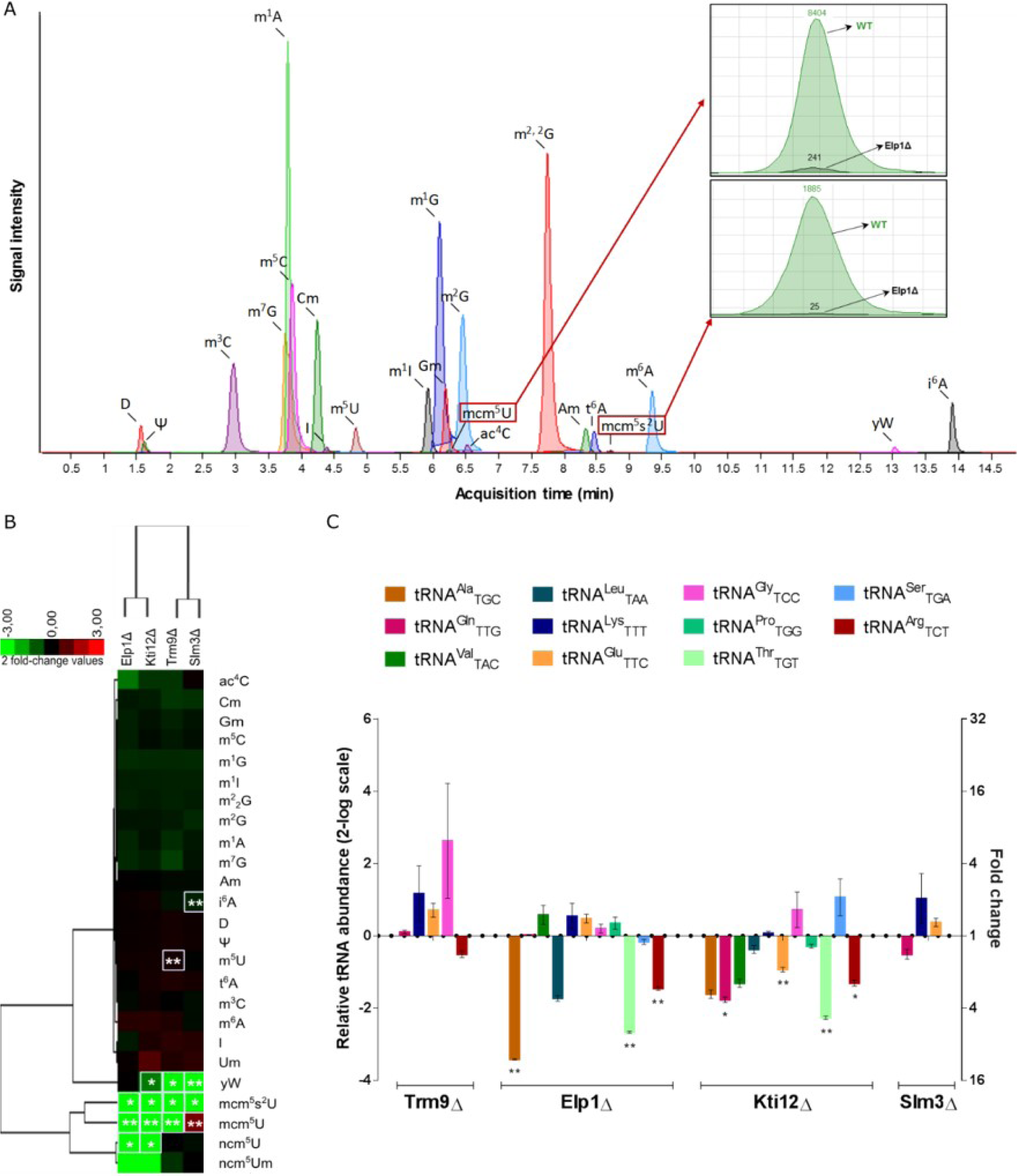
Knocking out tRNA-modifying enzymes affects steady-state tRNA modification profiles and abundance. **A**. Total ion chromatogram of LC-MS/MS analysis of modified tRNA ribonucleosides (WT). Chromatograms for mcm^5^U and mcm^5^s^2^U comparing WT with Elp1Δ strains confirm reduced modification levels in the Elp1Δ strain. Data for other strains are shown in supplementary Figure S2. **B**. Hierarchical cluster of the relative abundance of tRNA ribonucleoside modifications in selected mutant strains. tRNA modifications were identified and quantified by mass-spectrometry. Log-transformed fold-change values for mutants were normalized against WT, and the resulting ratios were subjected to hierarchical cluster analysis. Data represent mean of triplicates of three independent clones (** p < 0.01, * p < 0.05, Student’s t-test with CI 95% relative to WT). Red: fold increase; Green: fold decrease; according to the scale in the top-left color bar. **C**. Quantification of wobble modified tRNAs in selected tRNAmod KO strains. The plot illustrates the abundance of each tRNA in mutant strains relative to quantified levels of the respective tRNAs in WT. Data was obtained using the four-leaf clover qRT-PCR method (37), and represents relative tRNA abundance ± SD in a 2-log scale of triplicates of 3 independent clones (calculated with REST software (38)).

Differences in detectable tRNA modification levels can result from either differential enzymatic activity or differences in tRNA isoacceptor abundance. To distinguish factors contributing to variations in tRNA modification abundance, we also used FT-qRT-PCR to measure the abundance of tRNAs that were hypomodified in tRNA-modifying enzyme mutants relative to WT (37). Our analysis suggests that levels of tRNAs modified by Trm9 and Slm3, respectively, are not significantly affected by tRNA hypomodification (Figure 4C), which agrees with previous studies of Trm9Δ and the Tuc1Δ thiolase (11). In contrast, we observed significantly reduced tRNA levels in both Elp1Δ (tRNA^Ala^_ncm5UGC_, tRNA^Thr^ _ncm5UGU_ and tRNA^Arg^ _mcm5UCU_), and Kti12Δ (tRNA^Gln^ _mcm5UCU_, tRNA^Glu^ _mcm5UCU_, tRNA^Thr^ _mcm5UCU_and tRNA^Arg^ _mcm5UCU_), suggesting that Elp1 and Kti12 mediate tRNA modifications either directly or indirectly regulate tRNA stability (Figure 4C). In aggregate, these findings indicate that tRNA modifications are important not only for proteostasis, but also the abundance (and, ostensibly, stability) of certain tRNA isoacceptors.

### tRNA mutants generate increased levels of codon-biased protein aggregates

We next sought to investigate the mechanistic link between altered tRNA modification, isoacceptor abundance and increased protein aggregation observed in tRNA-modifying enzyme KOs. Using differential density centrifugation, we isolated aggregated protein fractions from exponentially growing WT, Trm9Δ, Elp1Δ, Kti12Δ, Slm3Δ and Elp3Δ cells. We used SDS-PAGE to visualize differences between total and insoluble protein fractions across mutants (Figure 5A and Figure S3), which confirmed the results of fluorescence aggregometry.

**Figure 5.**
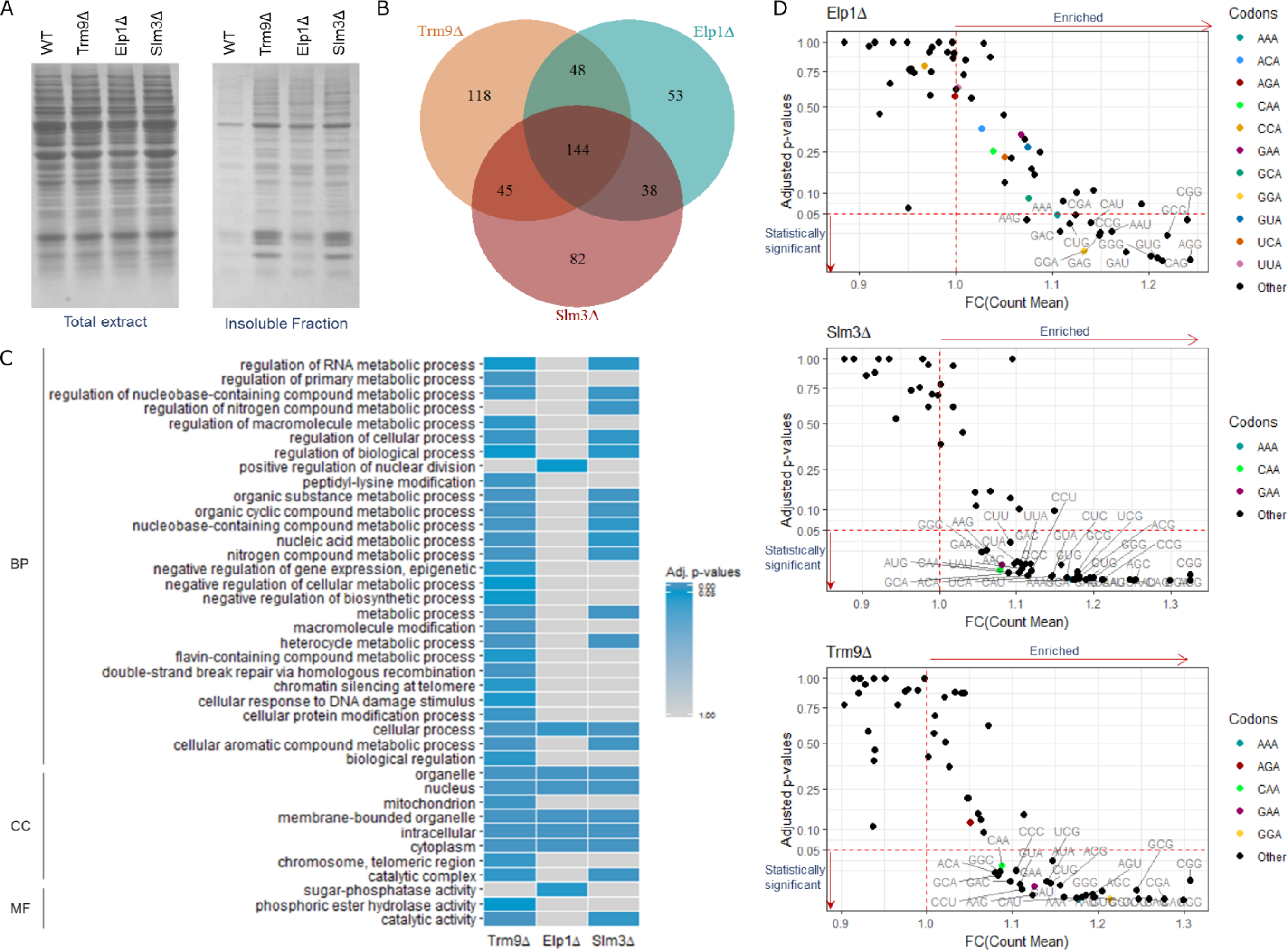
Mutant aggregomes exhibit significant functional enrichments and codon bias. **A**. SDS-PAGE (14%) and Coomassie staining was used to visualize total extracts and detergent-insoluble protein aggregates isolated using differential centrifugation of WT and mutants. **B**. Venn diagram of the overlap between upregulated aggregated proteins identified in selected mutants. **C**. Heatmap illustrating functional enrichment amongst upregulated proteins identified in selected mutants. Significant enrichments of biological processes amongst detected insoluble proteins were identified using GO data from https://go.princeton.edu/cgi-bin/GOTermFinder. p-values were adjusted with Bonferroni correction and FDR < 5%. BP: biological process; CC: cellular component; MF: molecular function. **D**. Aggregated proteins demonstrated significant enrichments in codons decoded by hypomodified tRNAs in mutant cells. Scatter plots show significance of codon usage fold-enrichments in Elp1Δ, Slm3Δ and Trm9Δ up-regulated proteins relative to reference genome. Significance was determined using the FSA package in R. Dunn (1964) Kruskal-Wallis test was applied for multiple comparisons and *p*-values were adjusted with the Benjamin-Hochberg method. Codons decoded by hypomodified tRNAs are colored. The red horizontal line indicates a *p*-value of 0.05 and the red vertical indicates a fold change of 1; codons below the horizontal line have a statistically significant enrichment and codons on the right of the vertical line are enriched relative to the reference genome. As in Trm9Δ and Slm3Δ strains, codons decoded by hypomodified tRNAs in Elp1Δ demonstrate general overrepresentation amongst aggregates, but these enrichments do not meet our criteria for statistical significance.

We then used LC-MS/MS to identify and quantify insoluble proteins from the mutants Elp1Δ, Slm3Δ and Trm9Δ. Relative to WT, we measured the significant upregulation (log_2_FC > 2) of 283, 309 and 355 proteins in Elp1Δ, Slm3Δ and Trm9Δ, respectively (Tables S7-S9). Among upregulated insoluble proteins identified in our analysis, 144 proteins were common to all mutant strains (Figure 5B).

We used gene ontology (GO) to analyze functional enrichments amongst aggregated protein fractions in mutant strains. This analysis revealed that aggregated proteins in Trm9Δ are involved in metabolic processes and their regulation, DNA repair, protein modification and regulation of gene expression (Figure 5C). Aggregates in the Elp1Δ strain were enriched in functions related to positive regulation of nuclear division and cellular processes, and Slm3Δ aggregates showed enrichment in functions related to metabolic processes and their regulation (Figure 5C). Collectively, GO results agree with the aberrant growth phenotypes observed in these mutants.

We hypothesized that genes encoding aggregated proteins may be enriched in codons decoded by hypomodified tRNAs in Trm9Δ, Elp1Δ and Slm3Δ (Tables 1 and S10), and used the Anaconda software plataform to analyze the full sequences of genes identified in our aggregated proteomic results for biased codon usage (49). Using functions of the Fish Stock Assessment (FSA) R package, we compared codon frequencies between mutant strains and the reference genome (S288c). This analysis identified significant enrichments of AAA, CAA and GAA codons (all of which are affected by mcm^5^s^2^U_34_) in Trm9Δ and Slm3Δ mutants; however, these enrichments did not extend to CAA and GAA codons in the Elp1Δ strain (Figure 5D). Previous studies have demonstrated that mcm^5^s^2^U_34_ and ncm^5^U_34_ modifications facilitate preferential decoding of both A- and, to a lesser extent, G-ending codons (50), while mcm^5^U_34_ promotes reading of both A- and G-ending codons, so we decided to analyze genes encoding precipitated proteins for overrepresentation of both A- and G-ending codons. As predicted, codons decoded by mcm^5^U_34_-modified tRNAs (e.g. AGG, GGA and GGG) were statistically enriched, relative to genome average, amongst genes identified from protein aggregates in Trm9Δ and Elp1Δ (Figure 5D). These results provide evidence that the loss of certain tRNA-modifying enzymes disrupts decoding at related codons and contributes to the precipitation of proteins encoded by codon-biased genes.

**Table 1.**
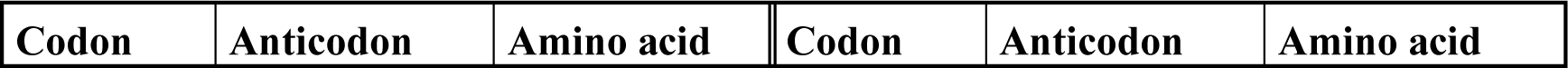
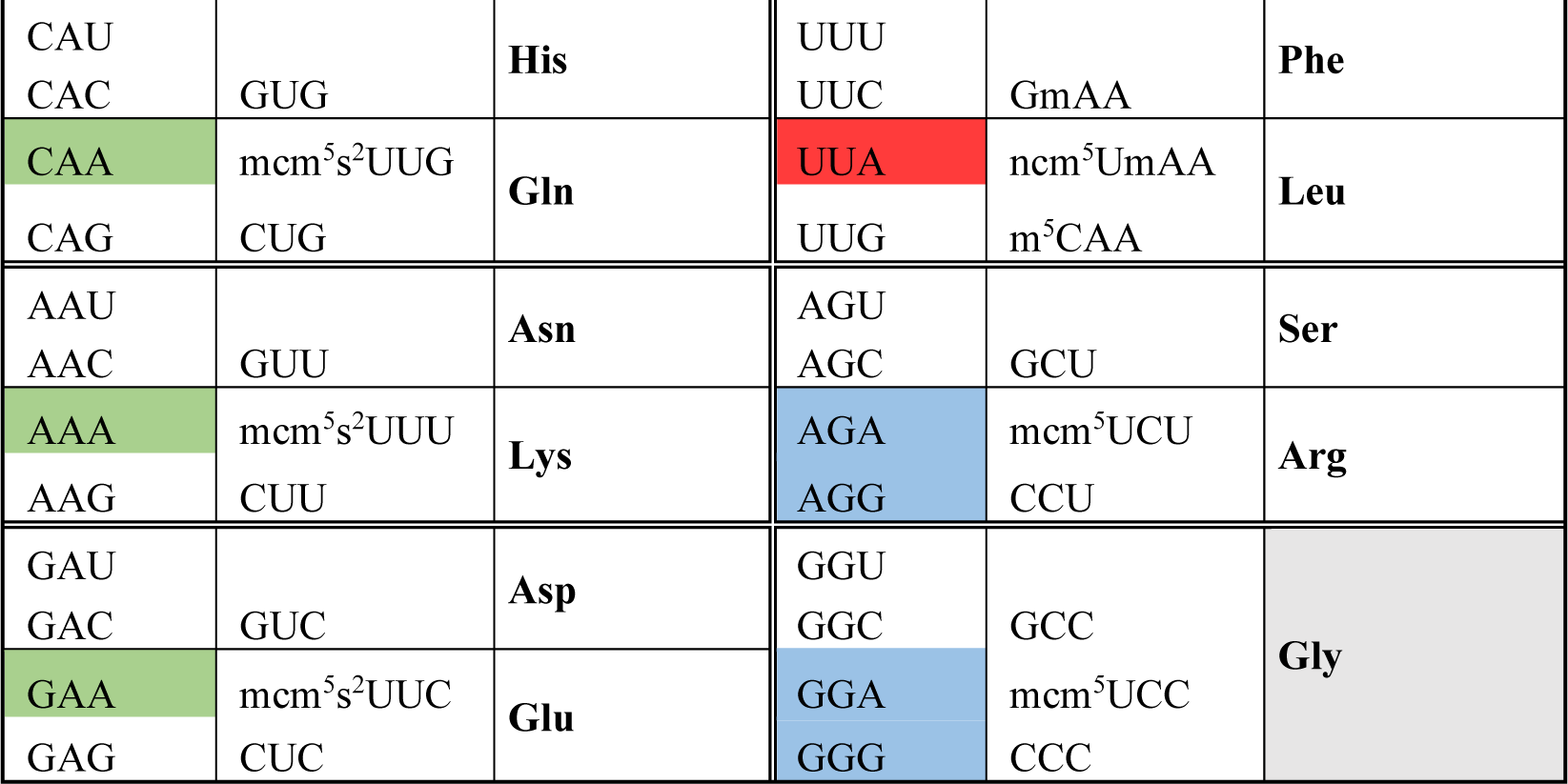
Codons whose decoding is affected by U_34_ hypomodification in two-split codon boxes. Loss of *slm3* perturbs translation of the 3 codons highlighted in green; absence of *trm9* affects the decoding of the 7 codons highlighted in blue and green; and deletion of *elp1* disrupts decoding of the 8 codons highlighted in red, blue and green.

### Loss of U_34_ modifications increases codon-specific mistranslation

We next sought to determine whether aggregated proteins were generated by increased translational termination in tRNA-modifying enzyme mutants, or by increased amino acid misincorporation caused by aberrant decoding of certain codons. Since ncm^5^U and mcm^5^s^2^U have been demonstrated to improve reading of A-ending codons, while mcm^5^U has been shown to facilitate decoding of both A- and G-ending codons (9, 11, 48, 50), and since wobble base modifications are generally known to prevent tRNA binding to U- and C-ending codons in two-split codon boxes (51, 52), it may be expected that mutants with hypomodified tRNA wobble bases experience higher rates of mistranslation. Because mcm^5^s^2^U_34,_ mcm^5^U_34_ and ncm^5^U_34_ are believed to enhance discrimination between cognate and near-cognate codons in mRNA, we hypothesized that mutants lacking these modifications experience higher rates of amino acid misincorporation due to expanded decoding of U- and C-ending codons. For example, loss of these modifications could permit decoding of AAA and AAG in place of AAU and AAC codons, and we predicted that misincorporations would be increased during translation of codons belonging to arginine (Ser_AGC_ to Arg_AGU_), glutamic acid (Asp_GAU_ to Glu_GAC_), glutamine (His_CAU_ to Gln_CAC_), lysine (Asn_AAU_ to Lys_AAC_) and leucine (Phe_UUU_ to Leu_UUC_) in mutant strains (Tables 1 and S10).

Through computational analysis of our proteomic results, we identified amino acid misincorporations in insoluble protein fractions from both mutant and WT strains (Figures 6A and S4), and found statistically higher rates of misincorporation relative to WT for Asp-to-Glu in Slm3Δ and Asn-to-Lys in Elp1Δ (Figure 6B). Furthermore, with only a few exceptions, we identified predicted amino acid misincorporations at higher rates in aggregated proteins from mutant strains than WT control (Figure 6B). We also detected amino acid misincorporations in near-cognate codon sites, including increased levels of Asn-to-Lys (AAU/AAC) and Asp-to-Glu (GAC/GAU) conversions in all mutants, as well as other strain- and site-specific mistranslational events: His-to-Gln (CAU) misincorporations were higher in both Trm9Δ and Elp1Δ, while Ser-to-Arg (AGC) misincorporations were higher in only Trm9Δ (Table S11).

**Figure 6.**
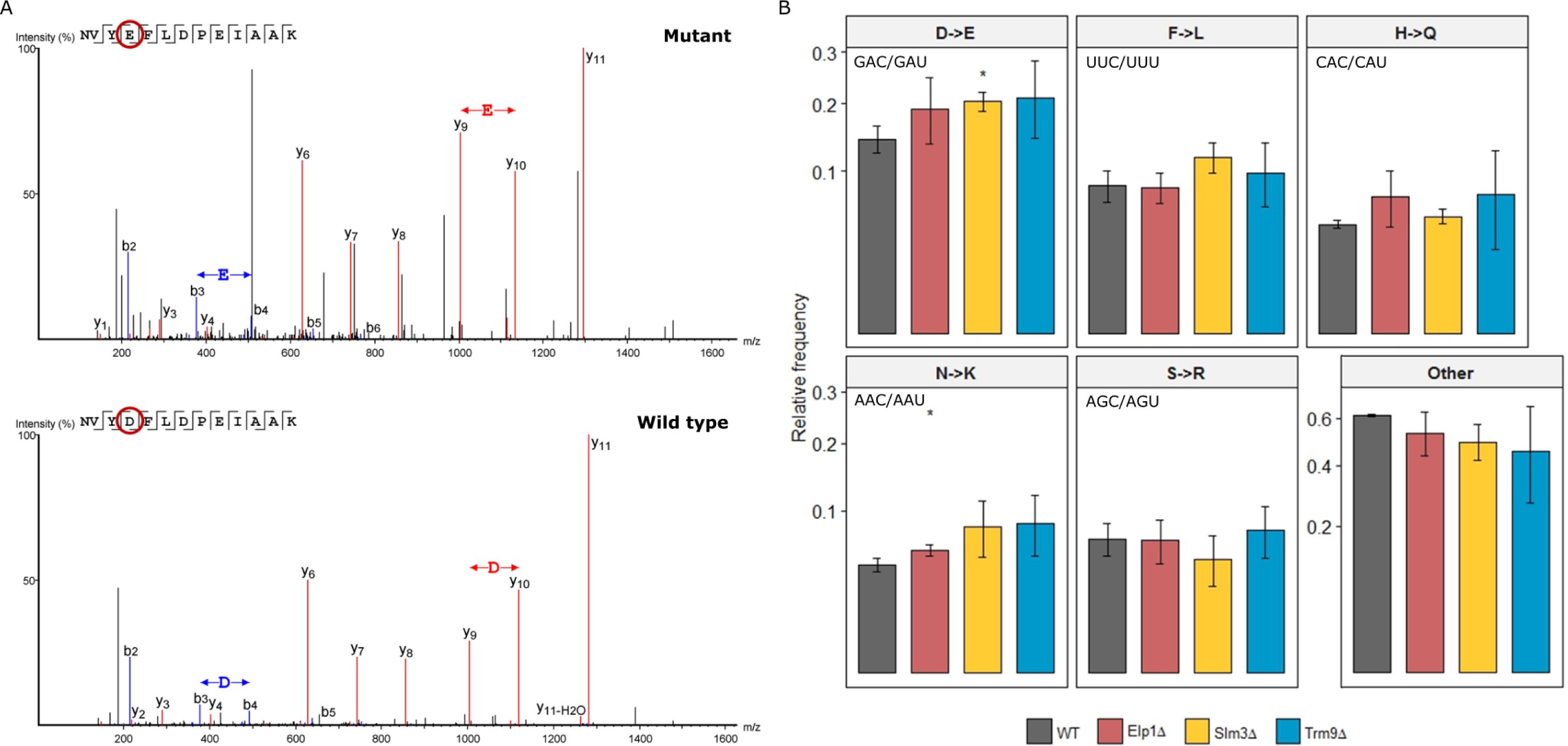
Aggregated proteins demonstrate increased rates of amino acid misincorporation in KO strains. **A**. Amino acid misincorporations were detected using SPIDER algorithm of the PEAKS Studio. Spectra exemplifying detection of one amino acid misincorporation of Glu at an Asp site (up spectrum) versus the WT peptide (bottom spectrum). **B**. Relative frequency of predicted and other amino acid misincorporations in mutant and WT strains.

We suspected that the absence of U_34_ modifications might also lead to protein misfolding and degradation by decreasing the decoding speed of a specific subset of codons overrepresented amongst the genes of aggregated proteins. To investigate this possibility, we analyzed the distribution of identified amino acid misincorporations across codons (Figure S5), and found that amino acid misincorporation occurred at higher levels at codon positions decoded by hypomodified tRNAs — particularly GCA and UUA in Elp1Δ, and AGA in Trm9Δ (Figure S5). Altogether, these findings underscore the critical importance of tRNA modifications for preserving translational fidelity.

## Discussion

While previous studies of single-gene deletions of tRNA-modifying enzymes have documented primarily subtle phenotypes, other reports have shown that disrupting more than one tRNA-modifying gene results in significantly altered cellular physiology, suggesting that epistasis may play a role in regulating tRNA modification profiles in yeast (32, 53, 54).

In this study, we engineered a genetic screen to assess the functional roles of 70 tRNA-modifying enzymes and measured increased protein aggregation in 5 strains. The sensitivity of our approach stems from the specificity of experimental techniques optimized for high-throughput fluorescence aggregometry; tRNA isolation, purification, modification profiling and isoacceptor quantification; and pan-proteomic detection of amino acid misincorporations.

Of the 42 cytosolic yeast tRNAs, 11 have modified uridine at position 34 — including ncm^5^U, ncm^5^Um, mcm^5^U and mcm^5^s^2^U (11). While at least 13 proteins are required for synthesis of ncm^5^ and mcm^5^ chains at U_34_ (55), our genetic screen showed that deletion of Elp1, Kti12, Trm9, Slm3 and Elp3 significantly altered protein aggregation (relative to WT: Elp1Δ: 4.9-fold; Kti12Δ: 5.1-fold; Trm9Δ: 3.4-fold change; : Slm3Δ: 3.1-fold; : Elp3Δ: 6.5-fold). While we measured significant decreases in U_34_ modifications in Elp1Δ, Kti12Δ and Trm9Δ strains (Figure 5B and Supplementary Figure 4), it remains curious that certain knockouts produced stronger protein aggregation phenotypes than others. Elp1 and its regulator Kti12 are critical for tRNA^Ala^ and tRNA^Thr^ ncm^5^U_34_ modification, and their deletion dramatically reduces the abundance of these tRNAs (Figure 5C). Our analysis also identified an association between U_34_ hypomodification and induction of specific RNA degradation pathways, which agrees with previous reports (53, 56–58). Interestingly, we found that the absence of a given tRNA modification (or its chemical derivatives) affected the abundance of at most two tRNA species, even though many tRNA modifications naturally occur in more than two tRNA isoacceptors. Similar observations have been reported in a survey of 20 sequenced yeast tRNAs lacking m^1^A_58_, a modification associated with drops in pre-tRNA^Met^_i_ levels (59). And the absence of m^7^G_46_ and m^5^C_49_ across 3 different tRNAs has been shown to result in the selective degradation of tRNA^Val^ _AAC_ (53).

Most amino acids are encoded by more than one codon (codon redundancy), however these codons are not used at the same frequency in all genes and influence gene expression in ways that remain incompletely understood. Growing evidence suggests that dynamic tRNA modification influences differential use of synonymous codons and may, therefore, regulate the selective translation of specific gene families (54). For example, independent studies have shown that modifications of the wobble position affect anticodon positioning in the ribosome, allowing for codon-dependent translation of specific transcripts, and that this mechanism is particularly relevant for the selective translation of DNA damage response genes (13, 51, 60, 61). Our analysis revealed that the aggregome of Trm9Δ yeast was enriched in proteins involved in the cellular response to DNA damage, which substantiates previous suggestions that loss of Trm9 may recapitulate the stress response to protein- and nuclei acid-damaging agents (60, 62).

Recent evidence suggests that tRNA activity is highly coordinated with mRNA codon demand (63), that some tRNA-modifying enzymes regulate protein expression by modulating translational speed at specific codons (64, 65), and that disrupting this decoding efficiency may lead to protein aggregation (66, 67). Furthermore, ribosome footprinting data demonstrated that non-modified U_34_ tRNAs bind poorly to the ribosome, reducing translational speed in a codon-specific manner. In particular, loss of mcm^5^ modification in Elongator-related mutants was shown to increase ribosome density at AAA, CAA and GAA codons located at the ribosomal A-site (68, 69), which agrees with our finding that aggregated proteins in Elp1Δ are enriched in AAA-biased genes (Figure 6D). This study also reported CAA and AAA codon enrichments at ribosomal A-sites (Ncs6Δ and Uba4Δ mutants) following loss of s^2^ (68), which again agrees with our findings that aggregated proteins in Slm3Δ are enriched in AAA-, CAA-, GAA- and GAG-biased genes (Figure 6D).

*In vitro* studies have shown that wobble base tRNA modifications play a role in translational fidelity (70) and molecular modeling, nuclear magnetic resonance and X-ray data showed that such modifications can alter the geometry of the ribosome-decoding center, promoting the binding of anticodons to their cognate codons (51, 52). mcm^5^U_34_ and mcm^5^s^2^U_34_ modifications have been implicated in differentiating between cognate and near cognate codons in split codon boxes and optimizing codon-anticodon interactions (42, 71). U_34_ hypomodification of glutamine, glutamic acid and lysine tRNAs reduced the speed of translation in a codon-dependent context (60, 72), a phenotype that was reversed by overexpression of the hypomodified tRNAs (46, 54). Two hypotheses sprang from these observations: (1) mcm^5^s^2^U_34_, mcm^5^U_34_ and ncm^5^U_34_ modifications enhance ribosomal binding of anticodons to cognate codons, increasing translational speed in a codon-dependent manner; and (2) the primary role of mcm^5^s^2^U and mcm^5^U modifications is not to reduce misreading of non-cognate codons ending with U or C in the split codon boxes, but rather to improve the reading efficiency of cognate codons ending with A. Since U_34_ modification defects lead to codon-specific translational pausing (67, 68), we propose that pausing affects co-translational protein folding (leading to aggregation of misfolded proteins), while also increasing the rate of amino acid misincorporations we report in the present study.

Hypomodified tRNAs of the Elp1Δ and Elp3Δ strains should read 4 codons by base pairing with all 4 nucleotides at the third position of codons and likely with near-cognate codons that belong to split codon boxes. Previous ribosome occupancy studies showed increased occupancy of only CAC and CAU codons in Elongator mutants (67), while our analysis of protein aggregates in Elp1Δ revealed statistically significant enrichments of AAC, AAU, CAU, GAC and GAU (near-cognate codons that can base pair with the Elp1Δ-hypomodified tRNAs; Figure 6D and Table S11). This suggests that amino acid misincorporation at near-cognate codon sites may contribute to protein misfolding and aggregation.

In summary, our findings suggest that the deletion of certain tRNA modifying enzymes leads to the loss of tRNA modifications that play critical roles in translational dynamics — including both tRNA maturation and the accurate folding of full-length proteins. Future studies ribosome profiling studies along with the experimental tools we have developed should allow us to investigate mechanistically how codon-specific tRNA modifications, affect proteome stability.

## Acknowledgements

We thank Eduard Sabidó and Guadalupe Espadas-Garcia from UPF/CRG Proteomics Unit, and Amanda Del Rosario and Richard P. Schiavoni from MIT Koch Proteomics Core for assistance with proteomics; Rui Fernandes and Francisco Figueiredo from HEMS, IBMC, for technical assistance with transmission electron microscopy; Vera Afreixo and Vera Enes from iBiMED for the statistics support.

