## Supplementary Figures for "tRNA-modifying enzyme mutations induce codon-specific mistranslation and protein aggregation in yeast"

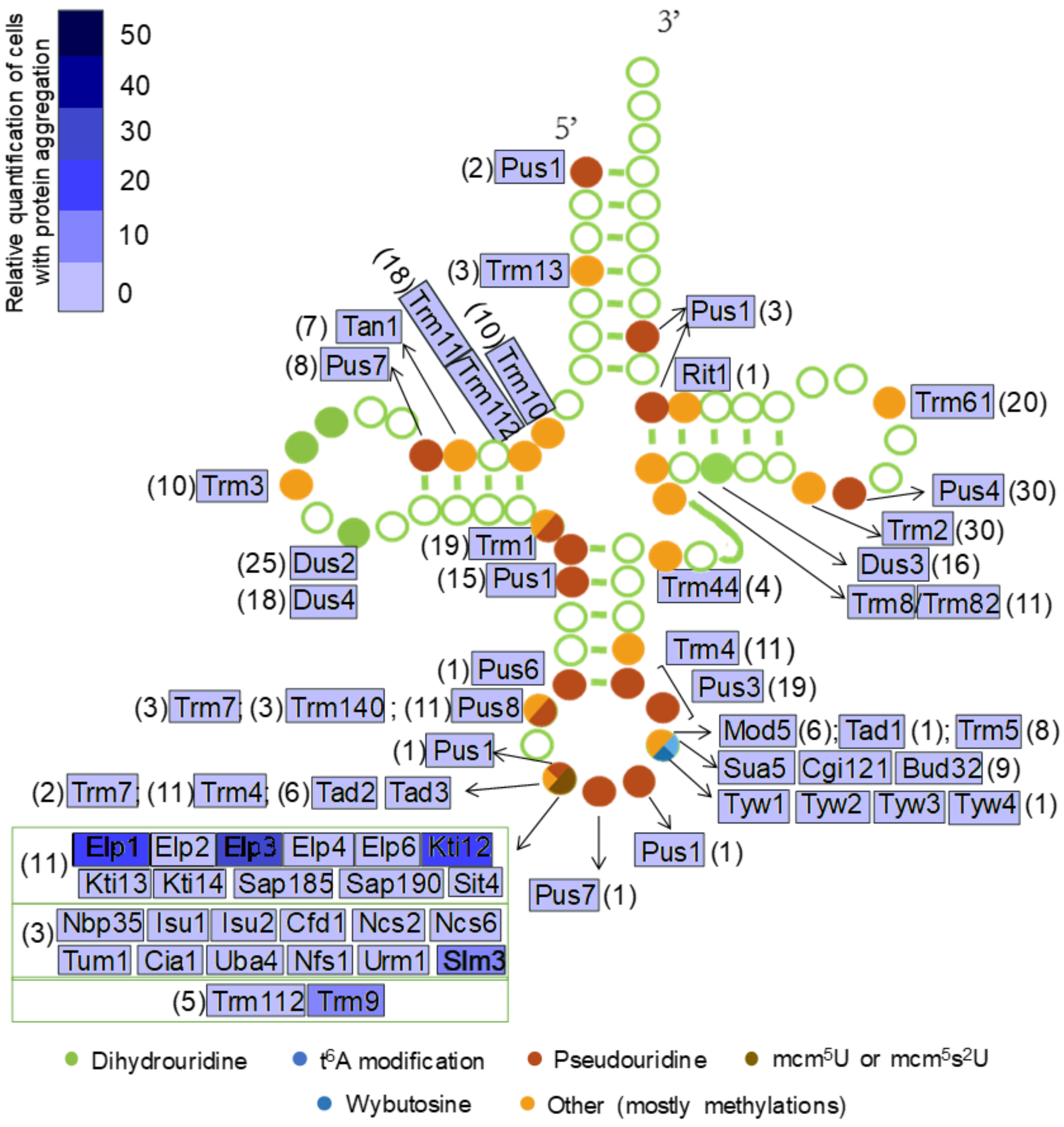


**Figure S1. Map of tRNA modifications catalyzed by enzymes investigated in this study.** The percentage of cells with fluorescent protein aggregates (average of three independent clones, relative to WT) is indicated for each tRNA modifying enzyme KO strain analyzed in this study. The number of different tRNA isoacceptors modified by each enzyme is also indicated in parentheses.

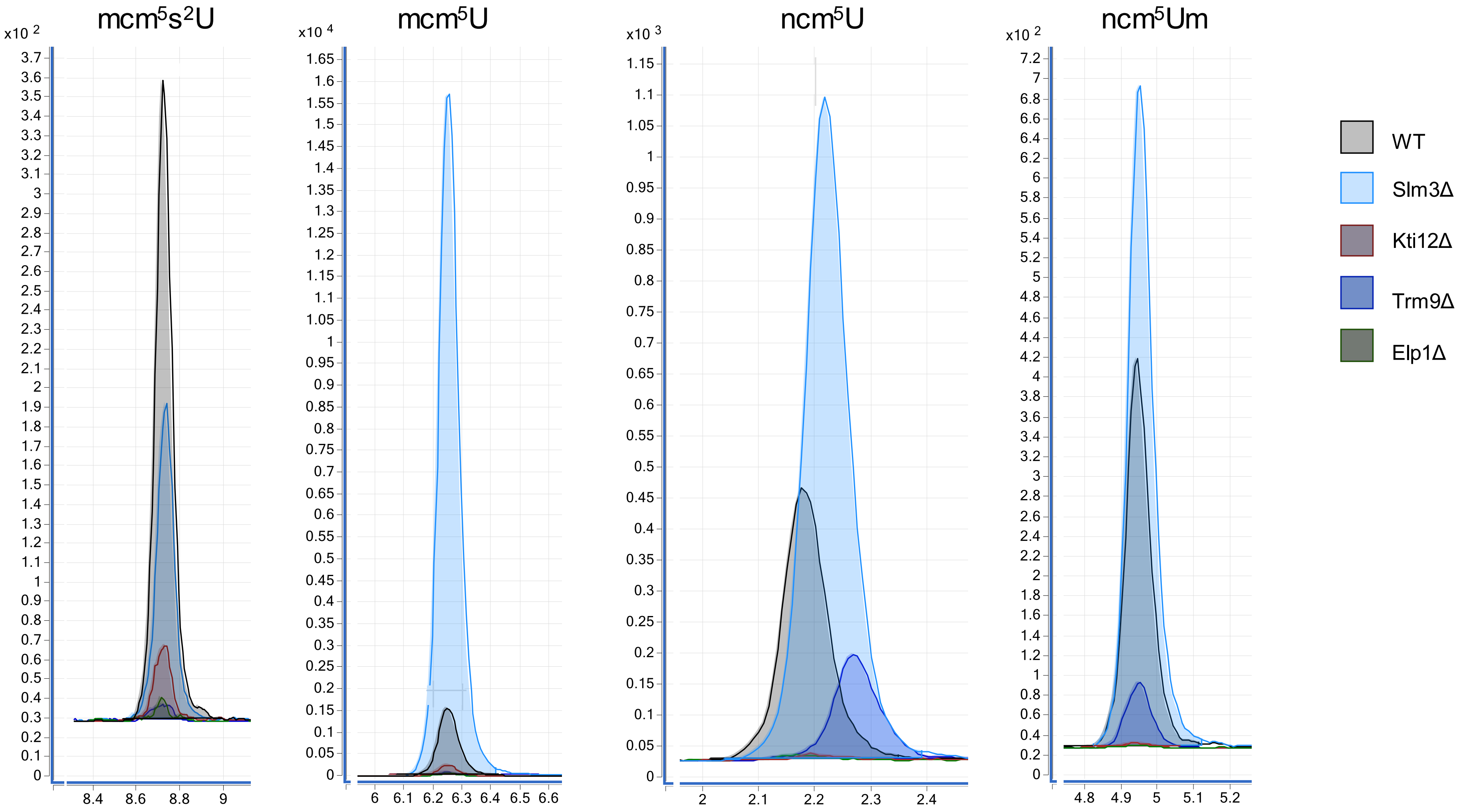
mcm$^5$s$^2$U
mcm$^5$U
ncm$^5$U
ncm$^5$Um
x10 2
x10 4
x10 3
x10 2
WT
Slm3Δ
Kti12Δ
Trm9Δ
Elp1Δ

**Figure S2. Knocking out tRNA-modifying enzymes reduces the abundance of tRNA wobble uridine modifications.** Chromatograms for $mcm^5s^2U$, $mcm^5U$, $ncm^5U$ and $mcm^5Um$ across WT, Elp1Δ, Kti12Δ and Trm9Δ strains confirms strain-specific suppression of wobble uridine modifications: (i) $ncm^5U$ and $ncm^5Um$ are absent in the Elp1Δ and Kti12Δ strains; while (ii) $mcm^5s^2U$ is decreased and $mcm^5U$, $ncm^5U$ and $ncm^5Um$ are increased in the Slm3Δ strain. Data for other modified nucleosides and statistical analysis are shown in supplementary Table S5.

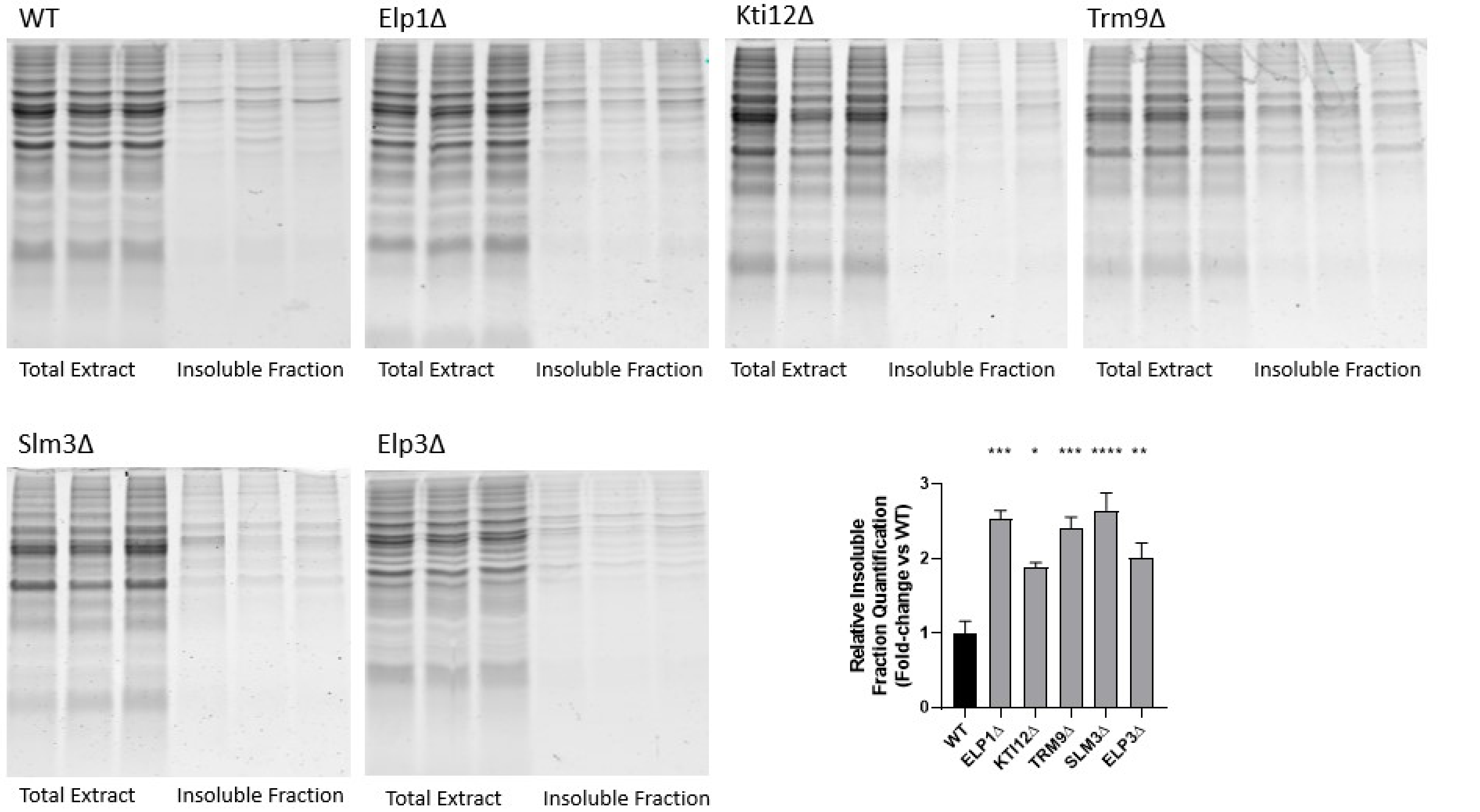


**Figure S3. Knocking out tRNA-modifying enzymes increases insoluble protein fractions.** Cells were grown to logarithmic phase in MM-His media at 30 °C and, after lysis, aggregated material was isolated, separated by SDS-PAGE and visualized by Coomassie staining. Gel lanes intensities were then quantified. Representative gel lanes showing the insoluble protein fraction of 3 clones of the selected KO and WT strains. The graph represents the relative quantification of insoluble proteins in selected KO strains relative to WT. Data represent the mean ± SD of three independent clones. (*<0.1; **p<0.01; ***p<0.001; p**** p<0.0001 One way ANOVA post Dunnett's multiple comparison test and CI 95% relative to WT)

A

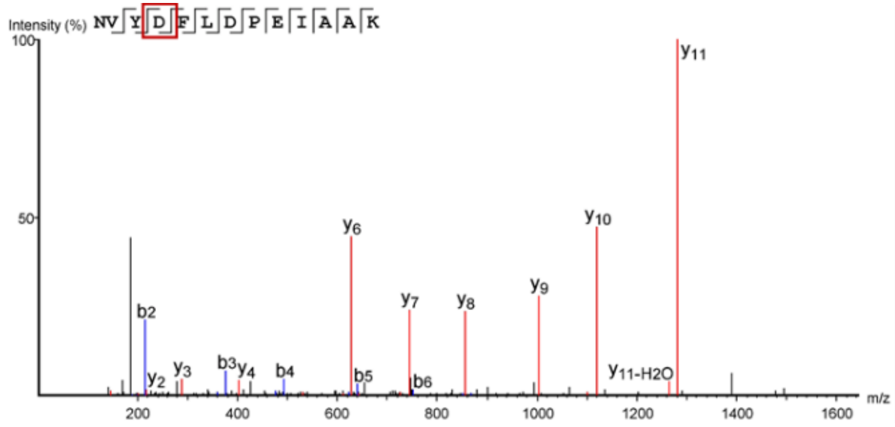


| # | b | b-H2O | b-NH3 | b (2+) | Seq | y | y-H2O | y-NH3 | y (2+) | # |
|---|---|---|---|---|---|---|---|---|---|---|
| 1 | 115.05 | 97.04 | 98.02 | 58.03 | N | | | | | 13 |
| 2 | 214.12 | 196.11 | 197.09 | 107.56 | V | 1380.70 | 1362.69 | 1363.68 | 690.85 | 12 |
| 3 | 377.18 | 359.17 | 360.15 | 189.09 | Y | 1281.63 | 1263.62 | 1264.61 | 641.32 | 11 |
| 4 | 492.21 | 474.20 | 475.18 | 246.60 | D | 1118.57 | 1100.56 | 1101.55 | 559.79 | 10 |
| 5 | 639.28 | 621.27 | 622.25 | 320.14 | F | 1003.54 | 985.54 | 986.52 | 502.27 | 9 |
| 6 | 752.36 | 734.35 | 735.33 | 376.68 | L | 856.48 | 838.47 | 839.45 | 428.74 | 8 |
| 7 | 867.39 | 849.38 | 850.36 | 434.19 | D | 743.39 | 725.38 | 726.37 | 372.20 | 7 |
| 8 | 964.44 | 946.43 | 947.41 | 482.72 | P | 628.36 | 610.36 | 611.34 | 314.68 | 6 |
| 9 | 1093.48 | 1075.47 | 1076.46 | 547.24 | E | 531.31 | 513.30 | 514.29 | 266.16 | 5 |
| 10 | 1206.57 | 1188.56 | 1189.54 | 603.78 | I | 402.27 | 384.26 | 385.24 | 201.64 | 4 |
| 11 | 1277.61 | 1259.59 | 1260.58 | 639.30 | A | 289.19 | 271.18 | 272.16 | 145.09 | 3 |
| 12 | 1348.64 | 1330.63 | 1331.62 | 674.82 | A | 218.15 | 200.14 | 201.12 | 109.57 | 2 |
| 13 | | | | | K | 147.11 | 129.10 | 130.09 | 74.06 | 1 |

B

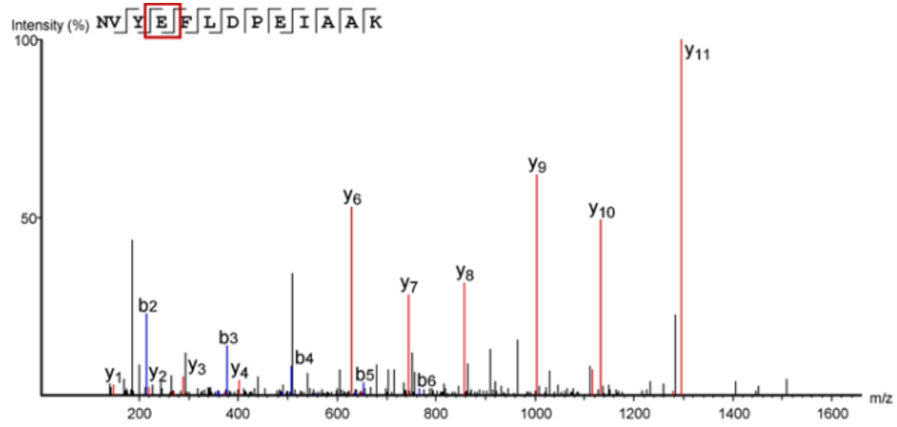


| # | b | b-H2O | b-NH3 | b (2+) | Seq | y | y-H2O | y-NH3 | y (2+) | # |
|---|---|---|---|---|---|---|---|---|---|---|
| 1 | 115.05 | 97.04 | 98.02 | 58.03 | N | | | | | 13 |
| 2 | 214.12 | 196.11 | 197.09 | 107.56 | V | 1394.72 | 1376.71 | 1377.69 | 697.86 | 12 |
| 3 | 377.18 | 359.17 | 360.15 | 189.09 | Y | 1295.65 | 1277.63 | 1278.62 | 648.33 | 11 |
| 4 | 506.22 | 488.21 | 489.20 | 253.61 | E | 1132.59 | 1114.58 | 1115.56 | 566.79 | 10 |
| 5 | 653.29 | 635.28 | 636.27 | 327.15 | F | 1003.54 | 985.54 | 986.52 | 502.27 | 9 |
| 6 | 766.38 | 748.37 | 749.35 | 383.69 | L | 856.48 | 838.47 | 839.45 | 428.74 | 8 |
| 7 | 881.40 | 863.39 | 864.38 | 441.20 | D | 743.39 | 725.38 | 726.37 | 372.19 | 7 |
| 8 | 978.46 | 960.45 | 961.43 | 489.73 | P | 628.37 | 610.36 | 611.34 | 314.68 | 6 |
| 9 | 1107.50 | 1089.49 | 1090.47 | 554.26 | E | 531.31 | 513.30 | 514.29 | 266.15 | 5 |
| 10 | 1220.58 | 1202.57 | 1203.56 | 610.79 | I | 402.27 | 384.26 | 385.24 | 201.64 | 4 |
| 11 | 1291.62 | 1273.61 | 1274.59 | 646.31 | A | 289.19 | 271.18 | 272.16 | 145.09 | 3 |
| 12 | 1362.66 | 1344.65 | 1345.63 | 681.83 | A | 218.15 | 200.14 | 201.12 | 109.57 | 2 |
| 13 | | | | | K | 147.11 | 129.10 | 130.09 | 74.06 | 1 |

**Figure S4. Computational detection of amino acid misincorporations in LC-MS/MS data.** The SPIDER algorithm of PEAKS Studio was used to identify single amino acid translational errors. Identified ions and corresponding amino acids are shown in tables below fragmentation spectra. **A.** Fragmentation spectrum from an accurately translated peptide. **B.** Fragmentation spectrum from a peptide with misincorporation of a single amino acid.

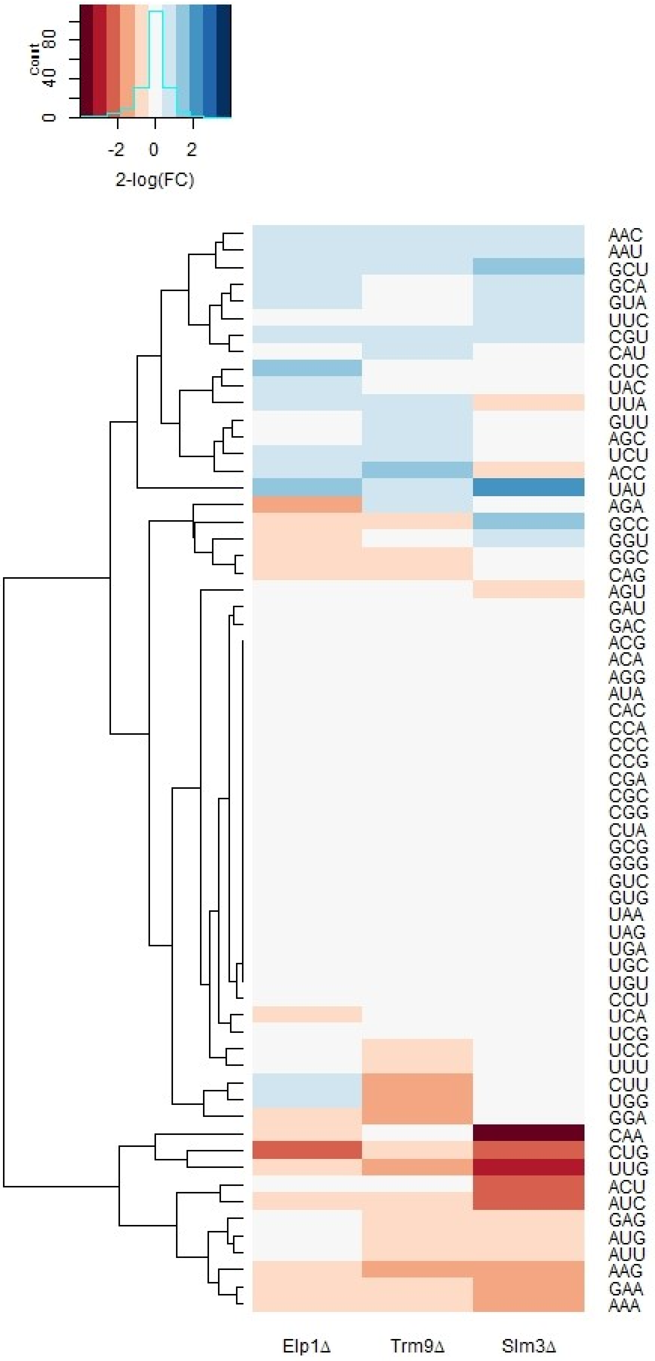


**Figure S5. Amino acid misincorporations localize in specific codon sites.** Amino acid misincorporations were identified using the SPIDER algorithm in PEAKS Studio, and an in-house R script was used to map detected translational errors to protein-coding genes. This heatmap illustrates the biased distribution of amino acid misincorporations in Elp1Δ, Trm9Δ and Slm3Δ (relative to WT).
